# Bill shape imposes biomechanical tradeoffs in cavity-excavating birds

**DOI:** 10.1101/2022.09.23.509155

**Authors:** Vaibhav Chhaya, Sushma Reddy, Anand Krishnan

**Affiliations:** Department of Biology, Indian Institute of Science Education and Research (IISER) Pune, Pashan Road, Pune 411008, India; Bell Museum of Natural History and Department of Fisheries, Wildlife and Conservation Biology, University of Minnesota, St. Paul, MN, USA; Department of Biological Sciences, Indian Institute of Science Education and Research (IISER) Bhopal, Bhauri 462066, Madhya Pradesh, India

**Keywords:** computed tomography, finite element analysis, barbets, beam theory, cavity excavation

## Abstract

Organisms are subject to a host of physical forces that influence morphological evolution. Birds, for instance, use their bills as implements to perform various functions, each exerting unique physical demands and selective influences on bill morphology. For example, birds that use their bills to excavate nesting or roosting cavities must resist a range of mechanical stresses to prevent fracture. However, the contribution of bill geometry and material composition to excavation stress resistance remains poorly understood. Here, we study the biomechanical consequences of bill diversification in two clades of cavity-excavating, frugivorous birds, the paleotropical barbets. Using multilayered finite element models and beam theory, we compare the excavation performance of different maxillary geometries for two loading regimes experienced by barbet bills during cavity excavation-dorsoventral impact and torsion. We find that deeper and wider maxillae perform better for impact loads than for torsional loads, with the converse for narrower maxillae. This results in a tradeoff between impact and torsion resistance imposed by bill geometry. Analytical beam models validate this prediction, showing that this relationship holds even when maxillae are simplified to solid elliptical beams. Finally, we find that composite bill structures broadly exhibit lower stresses than homogenous structures of the same geometry, indicating a functional synergy between the keratinous rhamphotheca and bony layers of the bill. Overall, our findings demonstrate the strong link between morphological evolution, behavior, and functional performance in organisms.

## INTRODUCTION

The evolution of biological form is, to a great extent, influenced by an organism’s interactions with its physical environment. Functional requirements of an organism, such as feeding and locomotion impose physical demands on biological structures, which in turn exert selective pressures on morphology (1, 2). Conversely, morphological diversification occurring under genetic and developmental constraints may influence functional performance, and organisms may adopt behavioral strategies that circumvent these constraints (3). Examining the interplay between form and function thus offers crucial insights into diverse evolutionary processes, and their consequences for the ecology and behavior of organisms. However, much remains to be understood about how the properties of the environment influence morphology, in particular during demanding tasks that generate significant and potentially damaging physical stresses.

The remarkable morphological diversity and multi-functionality of the avian bill renders it uniquely suited to address these biomechanical questions of form and function. Involved in numerous functions such as feeding, territorial defense and thermoregulation, bills are subject to a host of selective pressures on their shape and structure (4–8). In addition to these functions, many birds use their bills to manipulate substrate, from light to physically difficult maneuvers. Excavating nesting hollows in trees is a specialized behavior that is a particularly demanding task, because it requires the bill to withstand a range of compressive and shearing mechanical stresses in order to avoid structural failure (9). Consequently, cavity-excavating birds such as woodpeckers possess a suite of morphological and anatomical features which putatively improve their resistance to stresses experienced during cavity excavation (9–13). However, the role of bill geometry in dissipating excavation stresses still remains poorly understood, which necessitates a comparative analysis of excavation performance across a range of bill shapes.

In addition to geometry, the bill’s response to mechanical loads is also determined by its material properties and composition. Bills possess a sandwich composite structure, with an outer keratinous sheath (the rhamphotheca) and an inner bony core that resembles a closed cell foam (14–16). This combination of materials confers the bill with an increased resistance to compressive buckling loads, ensuring a high strength-to-weight ratio (15). The contributions of this composite structural arrangement of the bill to cavity excavation also remain poorly understood. Comparative analyses of bill performance are challenging to undertake, in part because of the difficulty in obtaining live samples. However, a combination of computational approaches enables us to quantify both the effects of geometry and material properties on the stresses experienced during cavity excavation. Similar techniques have been used in a paleontological context, where live samples are impossible to obtain, and can provide important comparative insight into the relative effects of geometry on mechanical performance (17). These studies serve a predictive function, enabling us to draw inferences that can lead to testable behavioral and ecological hypotheses (17, 18). Finite element analysis (FEA) is a numerical method traditionally used in engineering to predict the mechanical response of complex structures to simulated loading conditions. The past two decades have seen an increase in the use of FEA to investigate the link between morphology and function in both extant and extinct taxa, with most studies primarily focused on feeding mechanics (19–24). For cavity excavating birds, a few studies have built finite element models of the woodpecker head to study the stress transmission patterns of the skull during impact (9, 12, 25, 26). However, comparative studies that use FEA to understand the contribution of bill shape to mechanical performance are lacking, and thus we also lack an understanding of how interspecific variations in geometry contribute to excavation performance.

Here, we investigate the consequences of both bill geometry and material composition on stresses experienced during simulated excavation in the paleotropical (Asian and African) barbets. These birds use their bills to carve out nesting or roosting cavities in dead or decaying branches, tree stumps, or termite mounds (27, 28). During the initial stages of cavity excavation, barbets typically employ two excavation techniques - a pecking action which involves an impact along the dorsoventral axis, and a gouging action which involves torsional loading along the longitudinal axis (27–30). The Asian and African barbets occupy different regions of bill shape morphospace, with African barbets generally possessing more convex maxillae with greater depth compared to the narrower and pointed maxillae of the Asian barbets (31). The diversity of bill shapes and excavation behaviors in these birds thus offers an excellent opportunity to comparatively explore the relationships between morphology, behavior and functional performance. We employ multilayered finite element models to address the following questions-a) How does bill shape and the loads experienced during different kinds of excavation behavior (impact versus torsion) affect functional performance, as measured by the relative stresses experienced during simulated excavation? b) Does the composite structural arrangement of the bill improve its resistance to excavation stresses? We validate the predictions from our finite element analysis by comparing them to predictions from classical beam theory, by approximating the complex structures of bills to solid elliptical beams. The theoretical bill performance space derived from beam models further enables us to ask-Can classical beam theory predict the relative performance of complex structures like the bird bill? By examining these questions, our study examines fundamental concepts underlying the biomechanical consequences of morphological diversification, and uncovers the links between evolutionary processes and mechanical function.

## MATERIALS AND METHODS

### CT data and segmentation

Our study performed a biomechanical analysis on the bills of 15 paleotropical Barbet species; 7 species belonging to the Asian family Megalaimidae and 8 species belonging to the African family Lybiidae. Each species chosen represented a major sub-clade within the two families, thus representing a synopsis of the morphological diversity in this group of birds (32, 33). Study skins for these species were loaned from the Field Museum of Natural History in Chicago and the American Museum of Natural History in New York (see Table 1 for a list of specimens). For each species, we chose the micro-computed tomography (μCT) scan of one specimen from a dataset that we collected for another study (refer to Chhaya et al. 2022 for a detailed description of scanning procedures) (31). Scans were selected on the basis of quality (intactness of the bill and absence of dirt or other extraneous material in the scans) for meshing, processing and simulation. Because much of the bill shape diversification in Asian and African barbets has been driven by changes in maxillary shape (31), we focused on studying the effect of maxillary geometry on performance during cavity excavation. To obtain the three-dimensional and multilayered structures of barbet maxillae for finite element analysis, we imported micro-CT scans into Amira 6 (ThermoFisher Scientific, Waltham, Massachusetts, USA) as DICOM image stacks, where we manually segmented them. First, we virtually dissected out the rhamphotheca anterior to the nares, by using the ‘labeling’ feature in Amira to classify voxels based on grayscale values. Next, we filled the rhamphotheca sheath with an endocast using the Magic Wand tool in each labeled slice. This endocast represented the inner bony core of the bill, with no gaps, as we required a manifold mesh for the finite element simulation. Finally, we used the SurfaceGen module in Amira to generate surface meshes for both these structures (rhamphotheca and inner bony core) and exported them as separate PLY files for further processing.

**Table 1.** List of specimens used for finite element analysis and beam models. FMNH- Field Museum of Natural History, AMNH- American Museum of Natural History.

| Species | Museum | Catalogue no. |
| --- | --- | --- |
| **Megalaimidae** |  |  |
| *Psilopogon haemacephalus* | FMNH | 218040 |
| *Psilopogon corvinus* | AMNH | 646655 |
| *Psilopogon javensis* | AMNH | 646702 |
| *Psilopogon viridis* | FMNH | 231405 |
| *Psilopogon lagrandieri* | FMNH | 76301 |
| *Psilopogon oorti* | AMNH | 646978 |
| *Caloramphus hayii* | FMNH | 303070 |
| **Lybiidae** |  |  |
| *Trachyphonus erythrocephalus* | FMNH | 113352 |
| *Trachyphonus purpuratus* | FMNH | 113364 |
| *Pogoniulus subsulphureus* | FMNH | 121846 |
| *Tricholaema leucomelas* | FMNH | 214390 |
| *Lybius leucocephalus* | FMNH | 113175 |
| *Lybius dubius* | FMNH | 278829 |
| *Stactolaema leucotis* | FMNH | 194917 |
| *Gymnobucco calvus* | FMNH | 185349 |

### Surface mesh processing

The PLY files for both the rhamphotheca and bony endocast were imported into Autodesk Meshmixer (Autodesk Inc., San Rafael, California, USA). To control for the effects of bill size, we used the Transform tool to scale the maxillae of all species to a length of 5 cm. This value was based on the length of the largest bills in the group, as a fixed reference point for analysis. Following scaling, we used the sculpting tools in Meshmixer to remove segmentation artifacts on the rhamphotheca, particularly in the zone of measurement (see below) for each species. At the end of the processing stages, we had scaled, clean meshes for the maxillae of all 15 species, which served as the input for simulations (Figure 1B).

**Figure 1.**
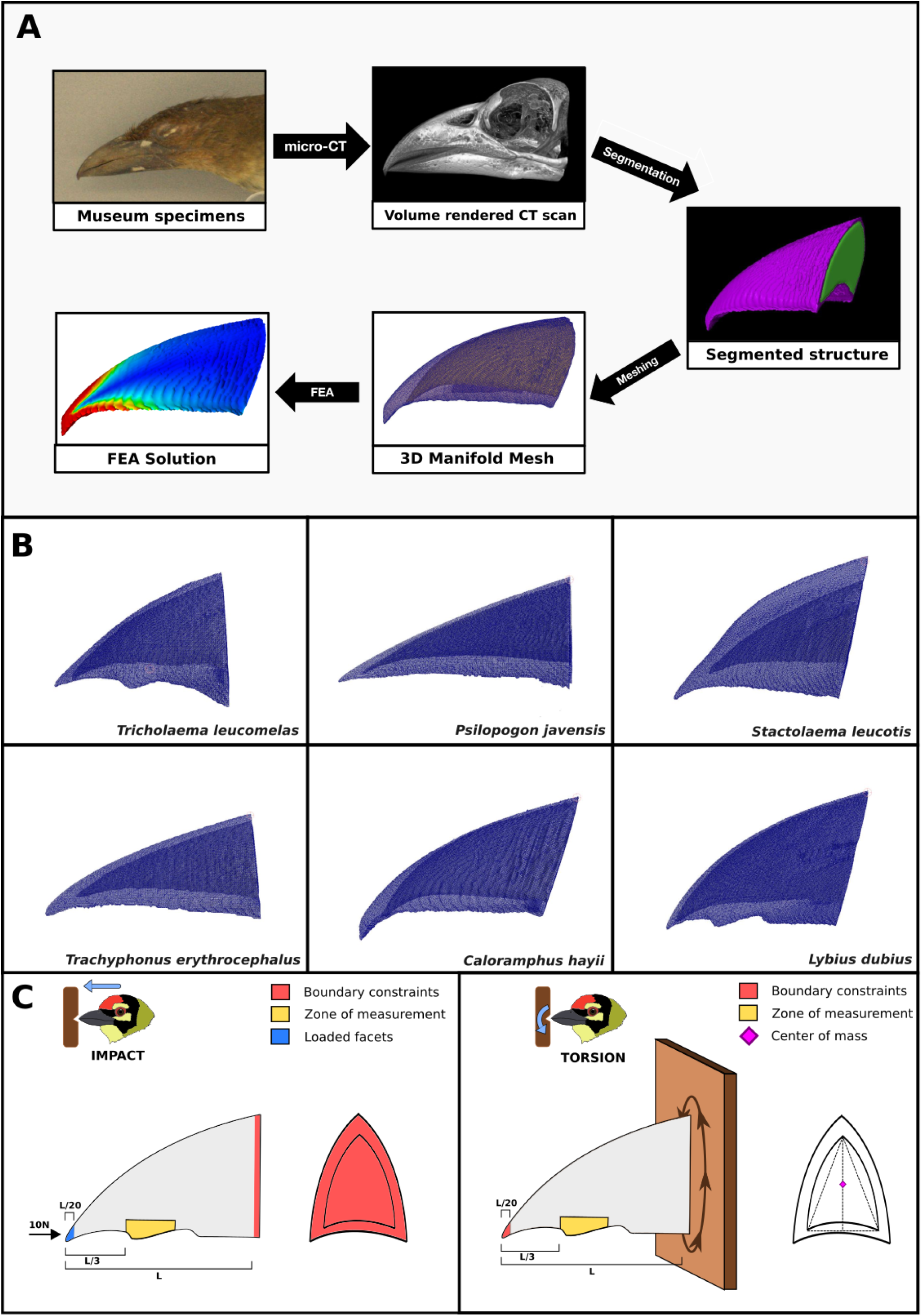
**A**. Workflow depicting the steps in the finite element analysis of barbet bills using museum skins, illustrated for a specimen of *Caloramphus hayii* (303070, FMNH). **B**. Manifold surface meshes for six exemplar barbet species, showing the outer rhamphotheca and inner bony core. **C**. Boundary conditions for the finite element models simulating impact (left) and torsion (right) along the dorsoventral axis.

### Finite element analysis

All finite element analyses were performed in FEBio Studio (Version 1.5.0) (34). Scaled and processed surface meshes of the rhamphotheca and bone endocast were imported and aligned to the world axes, with the tomial edge of the rhamphotheca extending anteriorly along the +*z* direction. For both structures, we generated Delaunay tetrahedral volume meshes with 4-node linear tetrahedral elements (Tet4) using the TetGen meshing tool in FEBio Studio.

We modeled the rhamphotheca and bony core as homogeneous, isotropic and linear elastic materials, with the values of material properties (Density, Young’s modulus and Poisson’s ratio) obtained from the available literature on bird bills. For Density (***ρ***) and Young’s modulus (***E***), we used values measured for the rhamphotheca (***ρ*** = 1000 kg/m^3^, ***E*** = 6.5 GPa) and trabecular bone (***ρ*** = 50 kg/m^3^, ***E*** = 12.7 GPa) of Toco Toucans (*Ramphastos toco*), which belong to the same superfamily (Ramphastoidea) as barbets (15). These values were measured using nanoindentation, which better represented our finite element simulations as homogeneous, isotropic materials. Because the Poisson’s ratio (***ν***) for avian bone and keratin has not been measured experimentally to our knowledge, we assigned a value of ***ν*** = 0.4 to both materials, following a recent study on Darwin’s finches which used these values (35–37). To compare the performance of a bone-keratin composite to a homogenous material composition of the bill, we also simulated two additional conditions-one in which both the rhamphotheca and endocast were assigned the material properties of keratin, and another in which both were assigned the properties of bone.

To connect the volume meshes of the rhamphotheca and bone, we specified a tied elastic contact between them, where the inner surface of the rhamphotheca was assigned as the primary surface and the outer surface of the bone endocast was assigned as the secondary surface. Barbets employ both pecking/chiseling actions and torsional gouging actions at the initial stages of cavity excavation (27–29). Based on this, we built dynamic finite element models for two loading regimes that barbet bills are likely to typically experience during cavity excavation-impact and torsion (Figure 1C). For the impact loading regime, we first fixed the nodes present on the basal surface of the rhamphotheca and bone meshes (no displacement along the ***x, y*** and ***z*** directions). Next, we applied a net force of 10N to the bill tip in the **− *z*** direction, on an area included within 5% of the tomial length (to ensure consistency across species and bill geometries). The chosen magnitude of force lies in the range of pecking forces experienced by the bill of the Great Spotted Woodpecker (*Dendrocopos major*), another primary cavity-excavator with a comparable body size to the barbets (9). Woodpeckers are close relatives of barbets, and thus provide a useful comparison as no direct measurements are available for barbets. For the torsion loading regime, we first fixed the nodes on the bill tip area that was loaded in the impact regime. Following this, we connected the base of the rhamphotheca and bone to a cuboidal rigid body, which had its area vectors aligned along the three world axes. Finally, we prescribed a positive rotation of ***θ*** = 1.75 milliradian (0.1°) to the rigid body parallel to the ***z*** axis, with the rotation axis passing through the center of the endocast base. This enabled us to compare the relative performance of different bill shapes across similar magnitudes of torsional force.

Using the Von Mises yield criterion, we quantified the excavation performance of barbet bills under these two loading regimes. We measured the peak Von Mises stress values at the outer surface of the rhamphotheca, in a zone present at two-thirds of the length of the tomium from the base (Figure 1C). This zone of measurement was selected such that it was away from the regions on which the boundary conditions were applied (the tip and base) in order to avoid local effects (20).

To ensure that mesh size did not affect the precision of our measurements, we performed convergence tests for the measured peak Von Mises stress on the best performing bills (lowest peak Von Mises stress) in each loading regime (*Tricholaema leucomelas* for impact and *Trachyphonus erythrocephalus* for torsion). From these convergence tests, we found that percentage error in VM stress falls below 5% on approximately doubling mesh size beyond 200,000 elements for both loading regimes (Supplementary data) (36–38). Thus, we ensured that all our species models had a mesh size greater than 200,000 elements for our finite element analysis.

### Beam models

Finally, to analytically validate the predictions made by our finite element models, we modeled barbet bills as solid elliptical beams of comparable dimensions undergoing axial loading and torsion. This analysis also served as an independent verification of how bill geometry influenced patterns of stress dissipation, by controlling for all other factors in a highly simplified mechanical model. We quantified the functional performance of these beams using two metrics analogous to the Von Mises stress values measured by finite element analysis in the impact and torsion loading regimes- critical buckling stress (***σ***_***cr***_) and maximum shear stress (***τ***_***max***_). These metrics were computed for each species as shown below (39).

Euler’s critical load, or the compressive axial load at which a slender column starts buckling about the ***x*** axis, is calculated by the formula

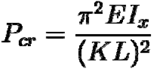

where ***E*** is the Young’s modulus of the column material, ***L*** is the length of the column, ***K*** is the effective length factor and ***I***_***x***_ is the second moment of area about the ***x*** axis. For a column that is fixed on one end and free to rotate and translate on the other (emulating the boundary constraints of the impact loading regime), the effective length factor ***K* = 2**. For an elliptical cross-section with principal axes ***a*** and ***b*** along the ***x*** and ***y*** axes respectively,

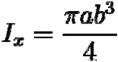

Thus, Euler’s critical load is given by

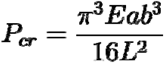

Dividing ***P***_***cr***_ by the area of the elliptical cross section ***A*= π*ab***, we get the critical buckling stress

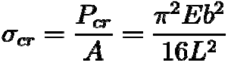

If a torque is applied to the free end of this elliptical beam (analogous to the torsion loading regime), the maximum shear stress (where higher stresses imply poorer performance) at the surface is given by

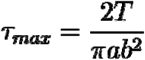

which occurs at ***x* = 0** and ***y = b***, for ***b* < *a*** where ***T*** is the resultant internal torque acting on the cross-section. For a given angle of twist ***θ*** and shear modulus of elasticity ***G***,

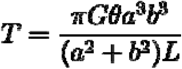

Thus,

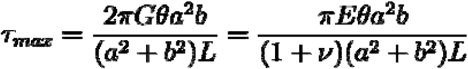

where ***ν*** is the Poisson’s ratio.

Using these formulae, we calculated the theoretical performance of elliptical beams for both axial loading (as an analogue of the forces experienced during impact) and torsion. We assigned the principal axes ***a*** and ***b*** of the elliptical cross-section as half of the bill width and depth respectively, measured at the base of the scaled rhamphotheca for each species using the measure tool in FEBio Studio. The length of all beams was set at ***L***=5 cm, in concordance with the standard bill length in our finite element models. Because the elliptical beams were modeled as homogeneous structures, we used the material properties of bill keratin (***E*** = 6.5 GPa, ***ν*** = 0.4) in our beam models. For torsion, we assigned a value of 1.75 milliradian to ***θ***, which was the angular displacement prescribed in the torsion loading regime.

To test if the theoretical metrics calculated from simplified beam models correlate with the Von Mises stress values from our finite element analysis for each loading regime, we performed Spearman’s correlation tests between them in RStudio (v1.3.959) (40).

## RESULTS

### Stress dissipation patterns of barbet maxillae

After calculating the von Mises (VM) stresses for all 15 species under both loading regimes, we mapped the distribution of these stresses over the surface of the bill. Heat maps depicting two example species (one each from Asia and Africa) are shown in Figure 2A. When bills were subjected to simulated impact, regions of high VM stress were primarily concentrated along the culmen (upper ridge) and tomium (lower edge), and dissipated longitudinally towards the base of the maxilla. VM stress dropped noticeably in the narrow region between the culmen and tomium, analogous to the neutral axis of a beam under a compressive axial load. When bills were subjected to simulated torsional loading, on the other hand, bills exhibited high VM stresses on the surface and on the region of the tomium near the tip, with a distinct null along the culmen instead. Thus, the two regions of peak VM stresses differed between the two loading regimes, indicating distinctly different patterns of stress propagation when the bill was used in tapping versus in a torsional gouging action.

**Figure 2.**
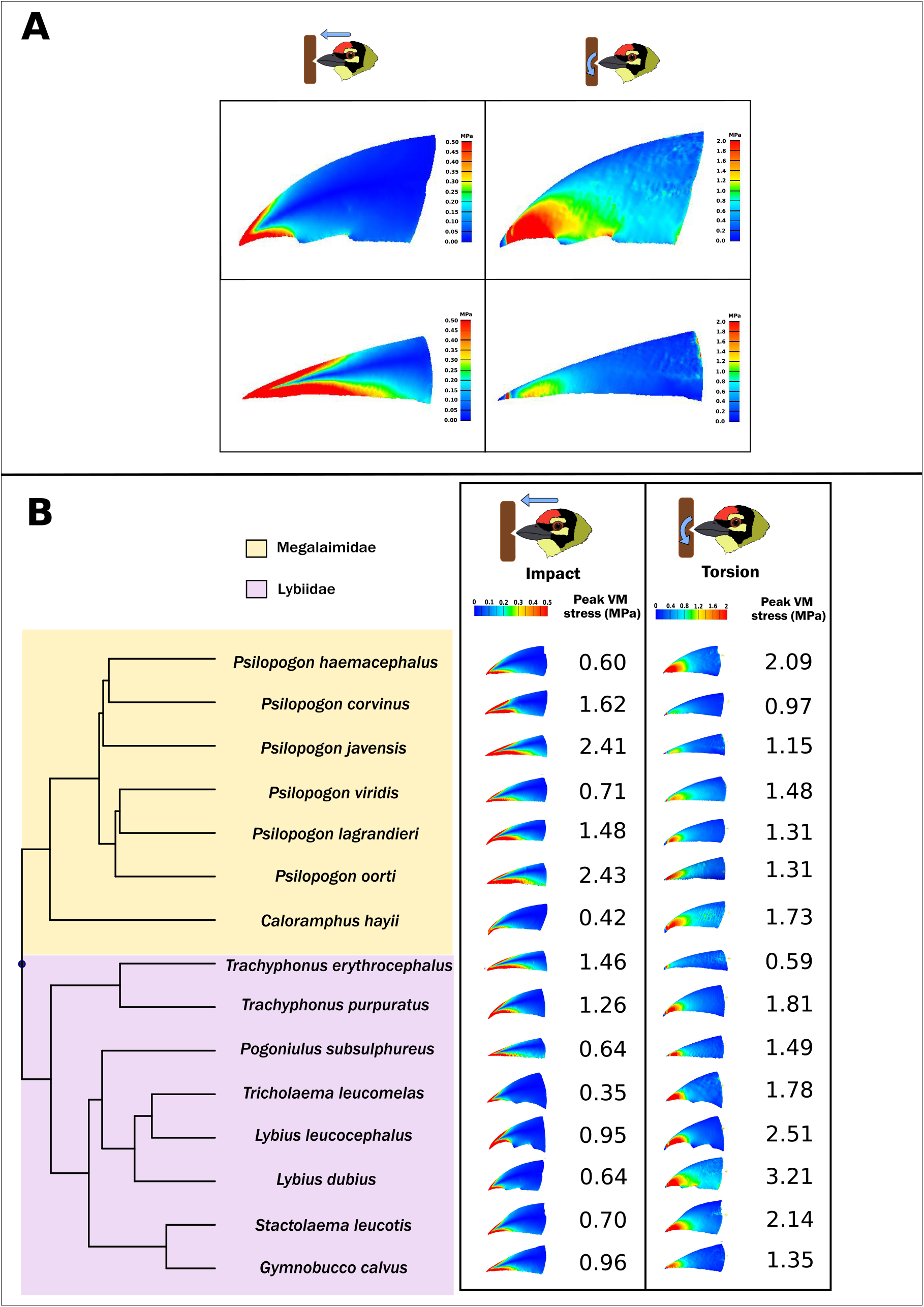
**A**. Distribution of Von Mises stresses on the maxillae of *Lybius dubius* (top row) and *Psilopogon javensis* (bottom row), for simulated impact (left column) and torsional (right column) loading. The deeper bill of *Lybius dubius* performs better for impact loading, while the narrower bill of *Psilopogon javensis* performs better for torsional loading. **B**. Distribution of Von Mises stresses and peak VM stress values for all species and loading regimes, mapped on to the pruned phylogeny of barbets (53).

### Bill shape and material properties influence stress dissipation patterns

We next sought to understand how bill geometry and material properties influence stress dissipation during excavation. To do this, we calculated the peak VM stress for all 15 species under both loading regimes (Figure 2B). Barbet species with relatively deep and wide maxillae, such as the African species *Lybius dubius* and *Tricholaema leucomelas*, typically exhibited comparatively lower peak VM stress values under simulated impact. In these bills, the VM stresses along the culmen and tomium dissipated at a relatively short distance from the tip. In contrast, species with maxillae that have relatively low width and depth such as the Asian species *Psilopogon javensis* and *Psilopogon oorti* exhibited higher peak VM stress values, and dissipated stresses relatively poorly along the longitudinal axis. On the other hand, when bills were subjected to simulated torsion, species with narrower bills outperformed those with deeper and wider bills. The relatively narrow-billed species discussed above all exhibited comparatively lower peak VM stress values for torsion, and this pattern was exhibited by both Asian and African species with different bill geometries. The peak VM stress values obtained for the two loading regimes were negatively correlated with each other (Spearman’s rho = -0.69, p = 0.006), further supporting a tradeoff between impact resistance and torsion resistance driven by bill geometry.

Next, we compared the peak VM stress between bills with homogenous material properties (outer layer and inner core both having the properties of either keratin or bone) and bills with a composite structure (keratin outer layer and inner bony core) (Figure 3). Under impact loading, composite structures performed better than homogeneous bone and keratin. The peak VM stress values measured for homogeneous structures were significantly higher (median percentage increase of 9.07% and 9.14% for bone and keratin respectively) (Wilcoxon signed-rank test, p<0.001 for both comparisons). For torsion, homogenous bone structures performed poorly compared to composite structures, showing a significant median percentage increase of 45.85% in the peak VM stress values (Wilcoxon signed-rank, p<0.001). Homogeneous keratin structures, however, performed marginally better than composites. Peak VM stress decreased compared to the composite (median percentage decrease of 5.80% in peak VM stress) (Wilcoxon signed-rank, p=0.006). On the whole, however, the composite bill structure, which represents natural conditions, exhibited better performance than homogeneous material across loading regimes.

**Figure 3.**
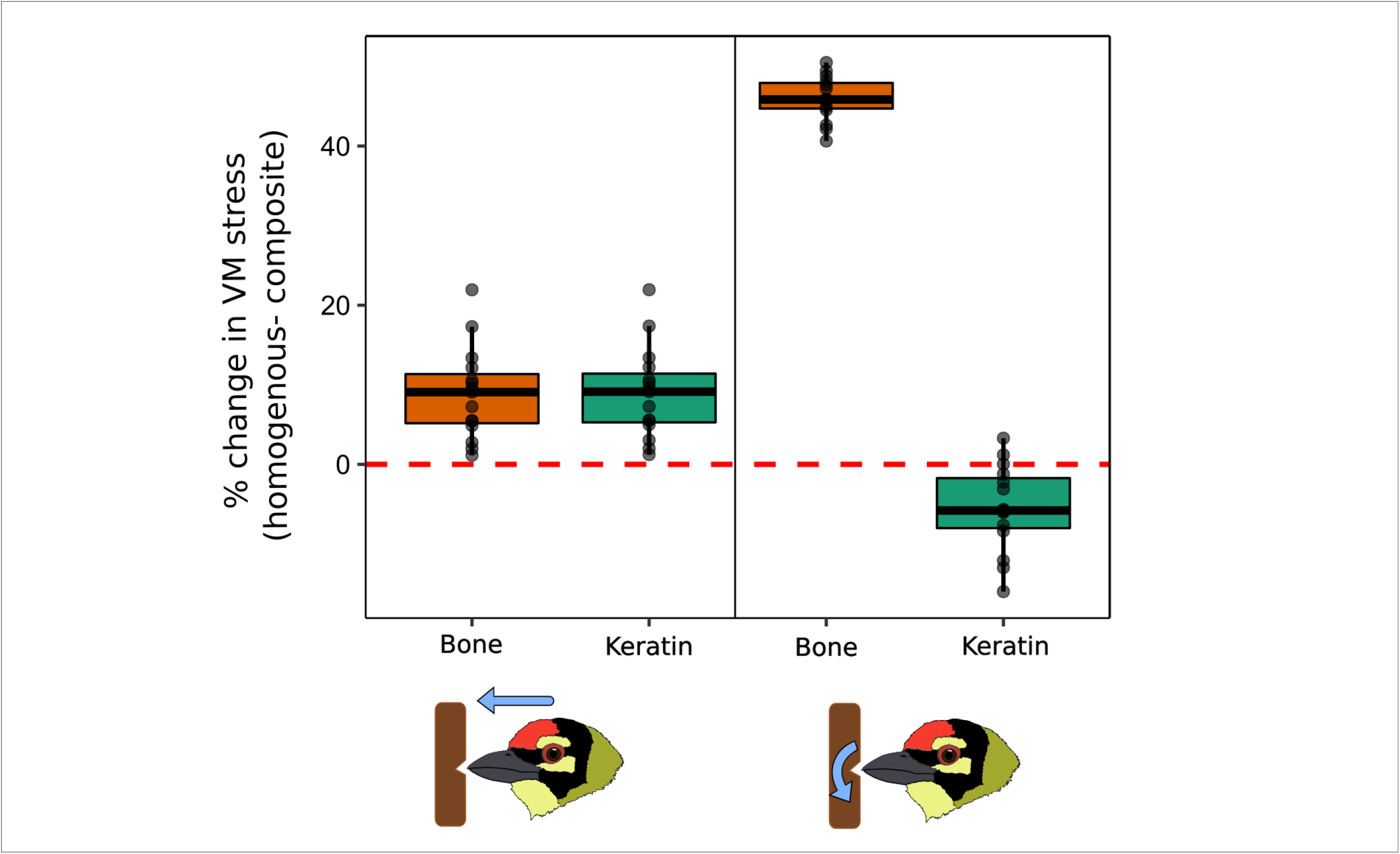
Boxplots depicting the percentage change in peak Von Mises stress values in homogenous keratin and bone compared to a composite structure, for impact (left) and torsion (right). The orange boxes represent the percentage difference in peak VM stress between the composite and a homogeneous bill made of bone, and the green boxes represent the difference from a homogeneous bill made of keratin. Values above zero indicate that the homogeneous material has higher stresses, and therefore that the composite performs better. Composites generally outperform homogeneous structures, as indicated by an increase in peak VM stress for homogeneous structures.

### A trade-off between impact and torsion resistance emerges from bill geometry

The results of finite element analysis suggested that barbet bills range from more impact-resistant to more torsion-resistant, putatively owing to variation in geometry. In order to quantitatively test this, as well as to validate the results of finite element analysis, we developed a simplified beam-theoretic model of the bill using dimensions obtained from the scaled three-dimensional structures of barbet maxillae. By simplifying the complex structures of barbet maxillae to solid elliptical beams, we predicted the effect of geometry on material failure for compressive axial loading (analogous to an impact) and torsion. According to the formulae outlined in the Methods, the critical buckling stress (***σ***_***cr***_), a measure of performance under conditions of impact loading, depends on bill depth. Therefore, deeper bills have a higher ***σ***_***cr***_, and perform better under compressive loads. We found that this prediction corresponded well with the results of our finite element analysis. Bills with higher depths performed better under impact loading (lower peak VM stress). The critical buckling stress (***σ***_***cr***_) exhibited a significant negative correlation with peak VM stresses for the impact loading regime (Spearman’s rho = - 0.70, p=0.006). The performance metric for torsional loading under our beam theoretic model, the maximum shear stress (***τ***_***max***_) depended on both depth and width of the elliptical beam. Thus, bills that are wider and deeper are predicted to exhibit high ***τ***_***max***_, and thus to perform poorly under torsional loading (see Methods). This is the inverse of the prediction under impact loading. Again, we observed a close correspondence between the theoretical performance space derived from beam theory with the measured peak VM stress values from our finite element models. Bills that were narrower tended to perform better under torsional loading. The maximum shear stress (***τ***_***max***_) is positively correlated with the peak VM stresses for the torsion loading regime (Spearman’s rho = 0.89, p<0.001). Thus, multiple methods independently verify that the biomechanical performance space of barbet bills represents a continuum between impact resistance and torsion resistance, with bill geometry as the primary driver of this tradeoff (Figure 4).

**Figure 4.**
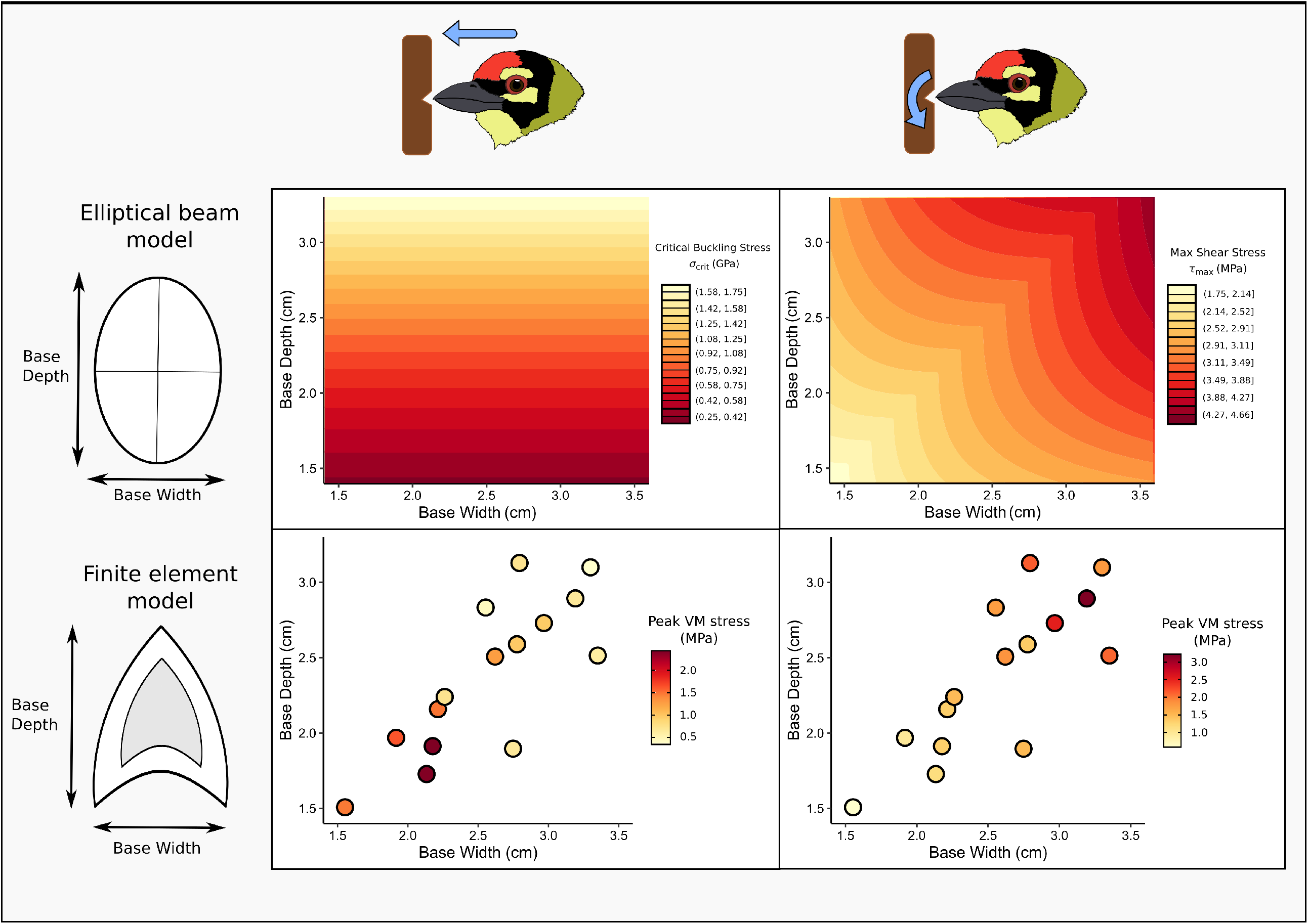
Excavation performance spaces obtained from finite element analysis (bottom row) and beam theory (top row), for impact (left column) and torsional (right column) loads. The bottom row plots the peak VM stress values (intensity of point color), against bill depth and width for all species. The top row plots the theoretical performance metrics (critical buckling stress for impact and maximum shear stress for torsion) calculated for elliptical beams of varying principal axes (corresponding to bill depth and width). Lighter colors indicate better performance. Both analyses show that bill depth and width are strong predictors of excavation performance, and impose a tradeoff between impact and torsion resistance.

## DISCUSSION

In summary, our comparative functional analysis reveals that bill geometry and material composition strongly influence cavity excavation performance in barbets. In our length-scaled finite element models, deeper and wider bills are more resistant to impact loading, whereas narrower bills exhibit higher resistance to torsional loading. Our analytical beam models further found that this relationship between bill geometry and functional performance was robust even when complex bill structures were simplified to solid elliptical beams, suggesting that stress dissipation by the bill is an outcome of simple geometric constraints. In addition to geometry, we found that the material composition of the bill, a sandwich-structured composite with an outer keratinous rhamphotheca and a low-density bony core, confers the bill with improved stress resistance compared to a homogeneous material composition. We note that our functional analysis serves to compare excavation performance across bill shapes and material compositions. Therefore, rather than identifying the actual failure limits for each model, our approach is comparative in nature, identifying which shapes better resist different loading regimes. Future kinematic studies including force measurements will help determine the actual failure points across the range of bill geometries, although the patterns we observe are likely to hold as they are validated by a number of robust theoretical analyses.

Performance tradeoffs, induced through mechanical and developmental constraints, have been commonly documented in biological structures (1, 3). A well-known example of this is the animal eye, which exhibits a tradeoff between sensitivity and resolution across species, as a result of structural and optical constraints on the eye (41). In geckos, the requirement for digital hyperextension during climbing results in a tradeoff between adhesion and running speed (42). The bird bill, as a multifunctional structure, is similarly subject to form-function tradeoffs. In Darwin’s finches, for example, bite force trades off with jaw closing velocity, therefore bills that specialize on more mechanically demanding food items are unable to produce song notes as quickly (43–45). Although the bills of barbets are also used to manipulate their diet of fruit, and may play a role in vocalization, cavity excavation is the most physically demanding task that the bill is used for. Because barbets eat soft fruit, and also vocalize with their mouths closed (27), the physical forces experienced during excavation are likely to pose a stronger selective constraint on bill geometry. Our study uncovers a mechanical tradeoff between impact and torsion resistance in the bills of cavity-excavating barbets. Beak depth and width were correlated with each other, constraining the range of bill geometries around a single axis in the morphospace (Figure 4). Our finite element and beam theoretic models both find that this axis represents a continuum, along which an increase in bill depth and width compromises torsion resistance in favor of greater impact resistance. Two testable hypotheses about the natural history of cavity excavation in barbets emerge from these theoretical predictions.

First, bill geometry may influence the excavation techniques employed by barbets, in order to minimize the stresses experienced by the bill during cavity excavation. Given their higher resistance to torsional loads, we expect barbet species with narrower bills (more commonly found in the Asian barbets) to employ gouging excavation behaviors, which involve torsion about the longitudinal axis. On the other hand, we predict that species with deeper and wider bills (more commonly found in the African barbets) should employ pecking behaviors during excavation, due to their higher resistance to dorsoventral impact. As a corollary to this, it is likely that all barbets employ a combination of both forms of excavation, but their relative use may depend on bill shape, as well as the substrate used for excavation, which leads to our second prediction.

Second, the geometry, excavation performance and behavior of barbet bills may be affected by the mechanical properties of available nesting substrates. African barbets, which occupy drier climatic regimes, exhibit deeper maxillae on average compared to the Asian barbets, which typically reside in more humid, forested regions (31, 32). Because climatic factors like temperature, precipitation and hydrological conditions affect wood decay rates and wood density, we surmise that they may also influence the availability of dead and decaying wood in the habitat (46, 47). These softer substrates are potentially easier to excavate due to their lower mechanical strength (48). Barbets may, therefore, be able to employ different strategies based on the availability of less demanding substrates. For example, *Trachyphonus erythrocephalus* exhibits a narrower bill with greater torsion resistance, and excavates holes in termite mounds, a softer substrate than wood (27). Thus, the mechanical properties of the bills may influence substrate selection by barbets. Additionally, differences in the availability of softer excavation substrates (49), may influence the physical demands placed on barbet bills during nest or roost site excavation, and consequently influence bill morphology. Cavity nesting is known to be a limiting factor on the distributions of birds that employ this technique (49–52), and our results suggest that biomechanical constraints could have ecological implications as well. Future studies incorporating mechanical testing of the substrate in various climatic conditions, and examining nest site selection by barbets in relation to the mechanical properties of this substrate, will help test these predictions.

From a materials perspective, we found that the composite structural arrangement of the bill generally performs better compared to a homogenous structure of the same geometry. This finding concurs with a study on the Toco Toucan (*Ramphastos toco*) bill, which found that the inner bony foam exerts a stabilizing effect on the outer rhamphotheca shell, providing greater resistance against deformation (15). A similar finding was reported in a study on the functional implications of edentulism in the therizinosaur *Erlikosaurus andrewsi*, where finite element analysis demonstrated that the presence of a keratinous rhamphotheca enhanced the stress dissipation ability of the skull for different biting scenarios (22). Thus, our study reveals a functional synergy underlying the integrated diversification of the rhamphotheca and bony modules in barbet bills (31).

At a broader level, our study demonstrates the power of comparative theoretical approaches such as finite element analysis and beam theory in quantitatively predicting the functional consequences of morphological diversification, particularly in poorly studied taxa such as barbets. By adopting a reverse engineering approach commonly employed in paleontological studies (17, 18), our study not only offers insights into the relationships between structure and function, but also serves as a launching point for future ecological and behavioral studies.

## ACKNOWLEDGMENTS AND FUNDING

We thank Nicholas Souza for scanning the barbet specimens at Loyola University Chicago, Thomas Sanger and James Cheverud for access to the scanner, Ben Marks, John Bates and Shannon Hackett at the Field Museum of Natural History, Chicago, Paul Sweet and Peter Capainolo at the American Museum of Natural History, New York. We additionally thank Sanjay P. Sane, Namrata Gundiah, Tanvi Deora, Yudhajit Kundu, and M.S. Madhusudhan for useful feedback and discussions. AK was funded by an INSPIRE Faculty Award from the Department of Science and Technology, Government of India, an Early Career Research Grant (ECR/2017/001527) from the Science and Engineering Research Board (SERB), Government of India and an initiation grant from IISER Bhopal. VC is the recipient of a KVPY Fellowship from the Government of India. SR was funded by a U.S. National Science Foundation grant DEB-1457624.

