## Supplementary data for "Bill shape imposes biomechanical tradeoffs in cavity-excavating birds"

**IMPACT-** ***Tricholaema leucomelas***

| S.No. | Elements | Facets loaded | Nodal force (kg cm/s^2) | Peak VM stress (kg/cm s^2) | % Error |
|---|---|---|---|---|---|
| 1 | 25644 | 33 | 30.3030303 | 4253.52051 | 0 |
| 2 | 63635 | 37 | 27.02702703 | 4100.6333 | 3.594369 |
| 3 | 131584 | 64 | 15.625 | 3643.14087 | 11.15663 |
| 4 | 240864 | 91 | 10.98901099 | 3495.49805 | 4.052625 |
| 5 | 565948 | 139 | 7.194244604 | 3325.97876 | 4.849646 |

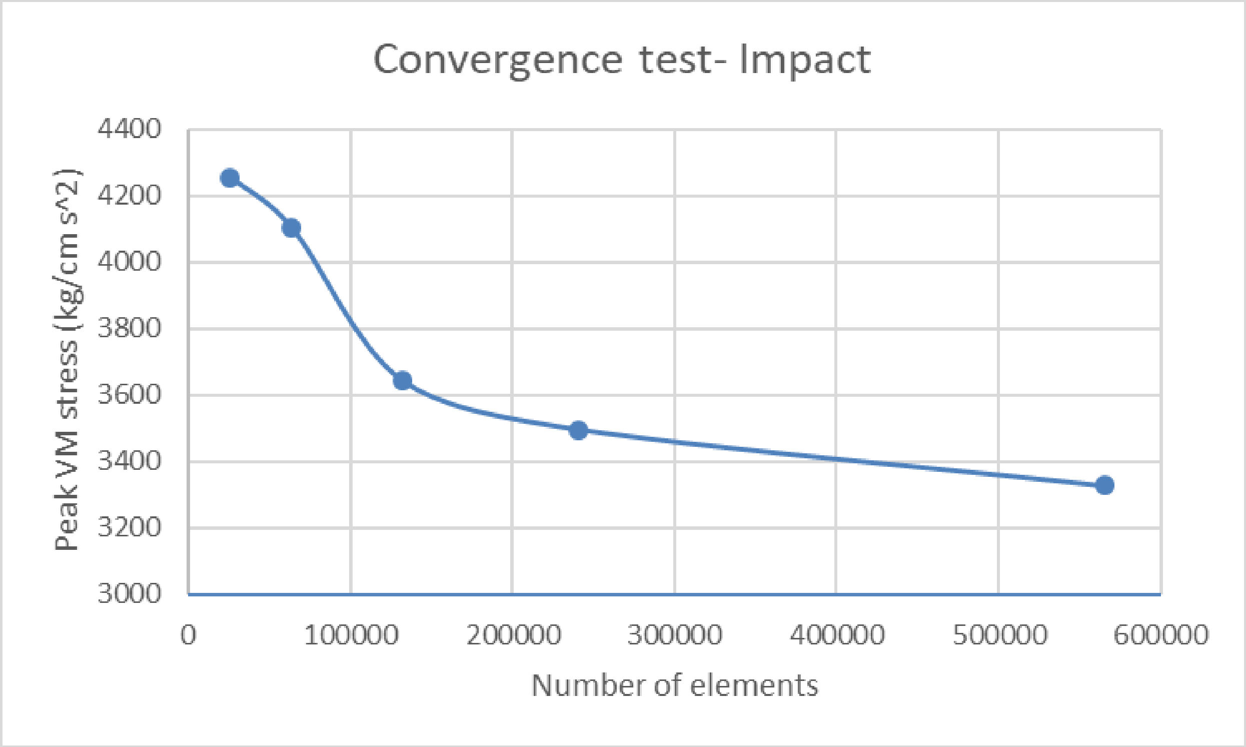

**TORSION- *Trachyphonus erythrocephalus***

| S.No. | Elements | Facets fixed | Peak VM stress (kg/cm s^2) | % Error |
|---|---|---|---|---|
| 1 | 35709 | 58 | 6275.38623 | |
| 2 | 63544 | 93 | 6252.29834 | 0.367912 |
| 3 | 152252 | 181 | 5606.03857 | 10.33636 |
| 4 | 263374 | 261 | 5843.979 | 4.244359 |
| 5 | 413006 | 382 | 5779.66309 | 1.10055 |
| 6 | 607507 | 504 | 5871.35059 | 1.586381 |

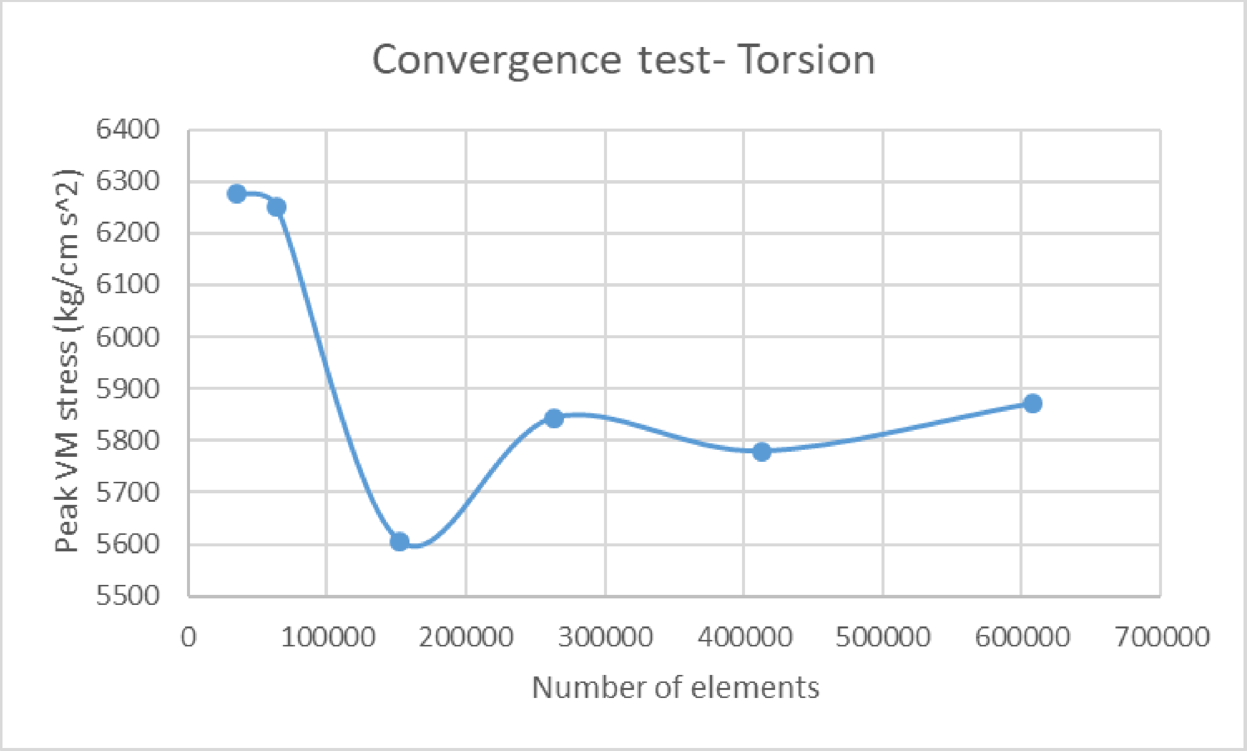
